# Protein and genetic interactions between RACK1A and FSD1 modulate plant development and stress granule-dependent response to salt in Arabidopsis

**DOI:** 10.1101/2025.02.25.640159

**Authors:** Pavol Melicher, Petr Dvořák, Maryna Tsinyk, Jan Řehák, Olga Šamajová, Kateřina Hlaváčková, Miroslav Ovečka, Jozef Šamaj, Tomáš Takáč

**Affiliations:** Department of Biotechnology, Faculty of Science, Palacký University Olomouc, Olomouc, Czech Republic

**Keywords:** superoxide dismutase, FSD1, RACK1, salt stress, stress granule, reactive oxygen species, antioxidant enzymes, development

## Abstract

The generation of reactive oxygen species (ROS) and their regulation by antioxidant enzymes such as IRON SUPEROXIDE DISMUTASE 1 (FSD1) are critical for managing plant responses to salt stress. However, the protein networks modulating ROS levels during salt stress remain incompletely understood. Our co-immunoprecipitation analysis identified the FSD1 as an interaction partner of the scaffolding protein RECEPTOR FOR ACTIVATED C KINASE 1A (RACK1A). Bimolecular fluorescence complementation analyses revealed that RACK1A interacts with FSD1 predominantly in the cytoplasm. Despite elevated FSD1 activity in *rack1a* mutants, the abundance of FSD1 protein remained unchanged. Computational predictions of interaction interfaces suggested that RACK1A may interfere with the catalytic site of FSD1. Advanced fluorescence microscopy and genetic studies further confirmed localization and relocation patterns of RACK1A and FSD1 during salt stress responses. Additionally, we found that the RACK1A-FSD1 module was involved in root hair tip growth, highlighting the developmental significance of this interaction. While *rack1a* mutants exhibited salt resilience, *fsd1-1 rack1a-1* double mutant displayed reduced salt stress resistance compared to *rack1a* single mutant, which was substantiated by deregulation of ROS levels. RACK1A and FSD1 accumulated in cycloheximide-sensitive structural condensates in the cytoplasm that colocalized with the stress granule marker protein TSN in roots. However, *RACK1A* knock-out completely abolished salt-stress-dependent relocation of FSD1 to structural condensates, suggesting that RACK1A recruits FSD1 to these stress granules. Thus, this study uncovers an entirely novel mechanism for the regulation of RACK1/FSD1-dependent antioxidant defense in response to salt stress in Arabidopsis.

## Introduction

Salt stress represents a severe constraint harming crop production (Hassani et al., 2021). Plants respond to salt stress by complex mechanisms governed by diverse signaling networks and second messengers, including reactive oxygen species (ROS) (Ovečka et al., 2014; Van Zelm et al., 2020; Singh, 2022). To regulate ROS levels, plants evolved a complex antioxidant system. Thus, gene modifications leading to antioxidant capacity reinforcement usually result in elevated salt stress tolerance (Zhang et al., 2014; Yan et al., 2016).

Superoxide dismutases (SODs) are key enzymatic antioxidants that catalyze the dismutation of superoxide into hydrogen peroxide (H_2_O_2_) (Pilon et al., 2011; Dreyer and Schippers, 2019). Recently, we demonstrated that the control of superoxide levels by FSD1 is inevitable for the efficient antioxidant defense during oxidative stress response (Dvořák et al., 2021; Melicher et al., 2022; Melicher et al., 2024). FSD1 is also involved in plant responses to salt stress, while FSD1 absence in *fsd1* mutants leads to a high accumulation of ROS and hypersensitivity (Dvořák et al., 2021). Nevertheless, molecular mechanisms regulating FSD1 compartmentalization and activity during salt stress are not entirely understood.

FSD1 and Cu/ZnSOD1 (CSD1) were identified by large-scale yeast two-hybrid screens as interactors of RECEPTOR FOR ACTIVATED C KINASE 1 A (RACK1A) (Guo et al., 2011). RACK1 is an evolutionary conserved cytosolic protein containing seven WD40 motifs forming a seven-bladed β-propeller structure that facilitates protein-protein interactions (PPIs) (Guo et al., 2019). Arabidopsis genome contains three *RACK1* isoforms: *RACK1A, RACK1B,* and *RACK1C* (Chen et al., 2006). *RACK1A* is a predominant isoform, as demonstrated by the pronounced phenotypes of the *rack1a* mutant (Guo and Chen, 2008). Plant RACK1 isoforms interact with proteins of diverse functions in plant development and responses to environmental cues (Islas-Flores et al., 2015; Masood et al., 2023; Li et al., 2023; Li et al., 2024). It might be involved in 60S and 80S ribosome assembly and translation by interaction with EUKARYOTIC INITIATION FACTOR 6 (Guo et al., 2011).

RACK1 is also implicated in salt stress response. Upon salt stress, UNIVERSAL STRESS PROTEIN 17 monomerizes and relocates to the nucleus, where it interacts with RACK1C, negatively affecting the expression of abscisic acid-mediated salt stress-responsive genes in Arabidopsis (Bhuria et al., 2022). Furthermore, overexpression of *Brassica oleracea RACK1* increased plant tolerance to the salt stress (Li et al., 2017), while *RACK1A*-overexpressing rice and soybean lines were hypersensitive to the salt stress (Zhang et al., 2018; Zheng et al., 2019). This suggests distinct roles of RACK1 in the salt stress tolerance of different plant species.

*RACK1* expression often correlates with the expression of several genes encoding antioxidant enzymes. In *Glycine max*, RNAi-mediated downregulation of *GmRACK1* was shown to positively affect *CAT, SOD, APX* and *GST* transcript levels and SOD, POD and CAT activities under salt and PEG-mediated drought stress (Zheng et al., 2019), while in rice, downregulation of *OsRACK1* reduced membrane peroxidation and enhanced SOD activity during drought stress (Li et al., 2009). Although the findings indicate the involvement of RACK1 in the regulation of antioxidant enzymes, the specific mechanisms of RACK1 contribution remain unclear.

The functional implications of RACK1A-FSD1 interaction may represent a key component of the plant antioxidant activity, but it remains largely unexplored. Here, we provide evidence that RACK1A interacts with and modulates the activity of FSD1. We revealed that RACK1A negatively regulates salt stress tolerance in Arabidopsis and accumulates in structural condensates identified as stress granules. In addition, RACK1A recruits FSD1 to stress granules, which likely contributes to the regulation of ROS levels during the salt stress response. Therefore, RACK1A plays a crucial role in the salt stress response in Arabidopsis, where the mechanism includes its interaction with FSD1 and relocation to stress granules.

## Material and methods

### Plant material and growth conditions

*Arabidopsis thaliana* (L.) Heynh ecotype Columbia (Col-0), referred to as wild-type (WT), *fsd1-1* and FSD1-GFP line (Dvořák et al., 2021), *rack1a-1* (Chen et al., 2006), *rack1a-5* (CRISPR-Cas9-mediated insertion mutant line prepared in this study) and *fsd1-1 rack1a-1* (prepared by conventional crossing in this study) mutant lines (all in the Col-0 background) have been used in experiments. We also generated transgenic *rack1a-1* bearing *5ˈUTR-pRACK1A::gRACK1A:GFP:3ˈUTR* (hereafter referred to as RACK1A-GFP line), which was transformed by floral dip method with *5ˈUTR-pFSD1::gFSD1:mRFP:3ˈUTR* construct. Additionally, the *5ˈUTR-pFSD1::gFSD1:GFP:3ˈUTR* construct from Dvořák et al. (2021) was transformed into the *rack1a-1* mutant. Selected lines with one insertion were propagated into the T3 homozygous generation.

Seeds were surface-sterilized by ethanol, dried and placed on a half-strength Murashige and Skoog (½ MS) medium and grown at 21°C and 70% humidity under a 16 h light/8 h darkness photoperiod with a photosynthetic photon flux of 120 μmol⋅m^−2^⋅s^−1^ in an environmental chamber (Weiss Technik, Grand Rapids, MI, United States) provided by cool white fluorescent linear tube light sources (Philips Master TL-D Reflex 36 W, light flow 3350 lm, light efficiency 93 lm⋅W^−1^) for a maximum of 14 days.

### CRISPR/Cas9-mediated RACK1A mutagenesis

We employed CRISPR/Cas9-mediated mutagenesis (Decaestecker et al., 2019) to prepare a knock-out (KO) *rack1a* mutant. Oligomers encoding *RACK1A*-specific guide-RNAs (gRNAs) were designed using CRISPR-P v2.0 (Liu et al., 2017) (Supplemental Table 1), synthesized and annealed. Entry (containing gRNA scaffold sequence, *AtU6* promoter sequence, and one of 4 annealed gRNAs) and destination vectors (containing *Ubi10::Cas9:mCherry*, *OLE1:mRuby* and all four *RACK1A*-specific *gRNAs*) were prepared by GoldenGate cloning (Engler et al., 2008). Expression vectors were introduced into *Agrobacterium tumefaciens* GW3101 and used for floral dip of WT plants. T1 generation seeds were selected by red fluorescence for the presence of *OLE1:mRuby* using a stereomicroscope (Axio Zoom.V16; Carl Zeiss, Germany). Random T2 generation plants expressing *Cas9:mCherry* were selected by stereomicroscope and gRNA1- or gRNA2-mediated mutations were detected by restriction fragment length polymorphism (RFLP) analysis in a 1793 bp sequence (containing all four gRNA complementary sequences) that was amplified using specific primers (Supplemental Table 2). Next, T3 generation plants not expressing *Cas9:mCherry* were selected by fluorescence microscopy and subjected to RFLP. The PCR products of selected plants were sequenced by Sanger sequencing using specific primers (Supplemental Table 2) and analyzed by DECODR (Bloh et al., 2021) and ICE CRISPR Analysis Tool (Conant et al., 2022) software. Finally, homozygous insertion mutant plants were selected in the T4 generation by sequencing.

### Preparation of transgenic rack1a-1 line complemented with GFP-tagged RACK1A under control of native promoter

Genomic DNA of *RACK1A* from WT, including native promoter sequence, *5ˈUTR* and *3ˈUTR* sequences, were used to prepare the fusion construct with a gene encoding *5ˈUTR-pRACK1A::gRACK1A:GFP:3ˈUTR* (referred to as RACK1A-GFP). Specific primers (Supplemental Table 2) were used to cover the sequence 1483 bp upstream of the start codon and 498 bp downstream of the stop codon. MultiSite Gateway® Three-Fragment Vector Construction (Thermo Fisher Scientific) kit was used to prepare the constructs. Briefly, PCR-amplified DNA of *5ˈUTR*, *RACK1A* gDNA and *3ˈUTR* were introduced into pDONR™P4-P1R (A fragment) or pDONR™P2R-P3 (C fragment) donor vectors by BP reaction to produce entry vectors. Constructs were introduced into *E. coli* TOP10 strain and positive clones were selected. Three-fragment LR recombination reaction was used to clone the fragments into the destination vector pB7m34GW. Positive clones were selected and confirmed by sequencing. T1 generation seeds were grown on a plant selection medium containing 10 μg·ml^−1^ phosphinothricin. Transgenic *rack1a-1* lines carrying RACK1A-GFP were propagated into T3 generation to ensure the presence of the transgene in both alleles.

### Phenotypic analyses of the mutant lines

Ten-day-old seedlings were documented for primary root length measurement using a flatbed scanner (ImageScanner III). The length of primary roots was measured by ImageJ (Schneider et al., 2012). Axio Zoom.V16 microscope was used to image root hairs and Zen Blue 2012 software (Carl Zeiss, Jena, Germany) was used to measure root hair length. The root phenotypic analysis was performed in three biological replicates. Fifteen plants per replicate were used for primary root length measurements. At least 300 root hairs from 20 plants were measured per replicate for root hair analysis. Mature root hair phenotypes were assessed in 7-day-old seedlings at a specifically designated distance from the primary root tip, ensuring measurements were taken more than 5 mm away. The fresh weight of 10-day-old seedlings was measured from 20 plants in three biological replicates for each plant line studied. Statistical significance was evaluated using a one-way ANOVA test in GraphPad Prism 8.3.0 (GraphPad Software, San Diego, California, USA).

### Immunoblotting and in-gel SOD activity analysis

To prepare protein extract, the powder of frozen plant material (100 mg) was resuspended and homogenized in 200 μl of extraction buffer (50 mM sodium phosphate (pH 7.8), 10% (v/v) glycerol, 2 mM ascorbate), placed on ice for 30 min and occasionally vortexed. The extract was centrifuged at 13 000 × g for 20 min at 4°C, and the protein concentration of the supernatant was measured according to the Bradford method (Bradford, 1976). The protein extract was used for the in-gel SOD activity assay. For immunoblot analysis, native protein extracts were supplemented with four times concentrated Laemmli SDS buffer at a 3:1 ratio and 5% (v/v) β-mercaptoethanol. The samples were boiled for 5 min at 95°C before SDS PAGE electrophoresis.

The immunoblotting and in-gel activity analyses were performed according to Melicher et al. (2022). Anti-FeSOD (AS06125) and anti-RACK1A (AS111810) primary antibodies were used for immunoblotting.

### Evaluation of salt stress responses

To determine the sensitivity of plants to salt stress, 4-day-old seedlings of WT, *fsd1-1*, *rack1a-1*, *rack1a-5*, *fsd1-1 rack1a-1* mutants and RACK1A-GFP-complemented *rack1a-1* mutant growing on ½ MS medium were transferred to the same medium containing 200 mM NaCl. The number of viable and fully bleached seedlings was counted on the 4^th^ day after transfer to NaCl-containing media and documented using a digital camera (NIKON Inc., Tokyo, Japan). Measurements were performed in three biological replicates with 30 plants per replicate.

### Light-sheet fluorescence microscopy (LSFM)

The plant material (RACK1A-GFP) preparation, mounting the seedling to fluorinated ethylene propylene (FEP) tube (Wolf-Technik, Germany) and its insertion into the observation chamber of the light-sheet microscope were performed as described previously by Ovečka et al. (2015). After 30 min of sample stabilization, developmental live cell imaging of the primary root was done by Zeiss Light-sheet 7 (Carl Zeiss, Germany) with dual-side illumination using two 10×/0.2 NA illumination objectives (Carl Zeiss, Germany) and pivot scan mode, excitation laser line 488 nm, beam splitter LP560, and emission filter BP505-545, detected with Plant-Apochromat 10×/0.5 numerical aperture (NA) water immersion objective (Carl Zeiss, Germany) and recorded by the PCO.Edge 4.2 camera (exposure time 20 ms). Time-course imaging was carried out in five consequential frames (Tiles) with an imaging frequency of every 1 min in Z-stack mode for 2-10 h. Developmental live cell imaging of lateral root formation was performed similarly, with single-side illumination using two 10×/0.2 NA illumination objectives and Plan-Apochromat 20×/1.0 NA water immersion detection objective (Carl Zeiss, Germany) with an exposure time of 100 ms and imaging frequency of every 10 min in the Z-stack mode for 12-17h. Stitching of tiles was done in Zeiss ZEN 3.4 (Blue version) software and data post-processing in Zeiss ZEN 3.4 (Blue version) or ArivisVision4D 3.0.1 software (Arivis AG, Rostock, Germany). Exported images were assembled into final figure plates in Microsoft PowerPoint.

### Confocal laser scanning microscopy (CLSM) of transgenic lines and image analysis

Live-cell imaging was performed using a CLSM on Zeiss LSM710 (Carl Zeiss, Germany), Zeiss LSM880 equipped with an Airyscan detector (ACLSM, Carl Zeiss, Germany) or Cell Observer SD Axio Observer Z1 spinning disk microscope (Carl Zeiss, Germany). Image acquisition was performed with Plan-Apochromat 20×/0.8 NA M27 (Carl Zeiss, Germany) and Plan-Apochromat 40×/1.4 NA Oil DIC (Carl Zeiss, Germany). A 488 nm excitation laser, set at 4% of the available intensity, was used for EGFP detection. Using CLSM, the emission of EGFP was detected in the 490 nm to 560 nm range. Emission ranging from 415 nm to 735 nm was used for ACLSM imaging of EGFP. For mRFP detection, a 561 nm excitation laser line was used, with emission detected between 590 nm and 650 nm.

To analyze localization changes in response to salt stress, seedlings of 4-day-old RACK1A-GFP line were placed in liquid ½ MS media between a glass slide and a cover slip. First, cells from the root differentiation and elongation zones were imaged to examine the RACK1A-GFP localization under normal conditions. To induce salt stress, 100 mM NaCl in ½ MS was added to the seedlings by perfusion. The RACK1A-GFP fluorescence was imaged for different time periods (30 min or 45 min). Recovery from salt stress was facilitated by replacing the solution with ½ MS medium without salt through perfusion. Cycloheximide (Sigma-Aldrich, C7698) treatment was performed using a 350 µM concentration (dissolved in distilled water) in liquid ½ MS medium with or without 100 mM NaCl.

ROS were visualized in root cells by incubating the samples in 30 μM CellROX Deep Red Reagent (Thermo Fisher Scientific), as described previously (Dvořák et al., 2021). The emitted fluorescence signal, excited at 639 nm, was captured within the 652–713 nm range using an Axio Observer.Z1 Spinning disk fluorescence microscope (Carl Zeiss, Germany). Images with CellROX Deep Red Reagent signal were processed as single planes using Zen Blue 2012 software (Carl Zeiss, Jena, Germany), exported and adjusted in Microsoft PowerPoint to final figures. The signal was analyzed using ImageJ software. Images were converted to an 8-bit grayscale format, and the mean signal density was measured.

Kymographs of growing root hairs were generated from time-lapse images acquired by LSFM along the line drawn longitudinally in the central part of growing root hairs from maximum intensity projections using the Zen software (Blue version). Semi-quantitative fluorescence signal intensity analysis was performed by a profile function in one central Z-stack of the selected compartments. Profiles for signal intensity distribution and quantitative evaluation were performed in Zen Blue 2014 software.

### BiFC assay – cloning and analysis

Arabidopsis total cDNA pool was used to amplify full-length cDNAs encoding *RACK1A, FSD1, NUDIX HYDROLASE 7* (*NUDT7*) and *GUANINE NUCLEOTIDE-BINDING PROTEIN ALPHA-1 SUBUNIT* (*GPA1*) using specific primers (Supplemental Table 2). Preparation of expression vectors was performed using “2in1” cloning system as described in Grefen and Blatt (2012). MultiSite Gateway® Three-Fragment Vector Construction (Thermo Fisher Scientific) kit was used to prepare the constructs. Expression vectors, carrying *RACK1A* fused to the N-terminal part of *YFP* (*YFPn*) as well as *FSD1*, *NUDT7*, or *GPA1* fused to the C-terminal part of *YFP* (*YFPc*) were prepared.

*N. benthamiana* leaf epidermal cells transiently transformed with *A. tumefaciens*, carrying the above mentioned vectors were used for semiquantitative rBiFC analysis. Confocal imaging was performed on Zeiss LSM710 (Carl Zeiss, Germany) with Plan-Apochromat 20×/0.8NA (Carl Zeiss, Germany) objective. A 514 nm excitation laser was used for YFP imaging, with emission detected in the 520 nm to 580 nm range. For mRFP detection, a 561 nm excitation laser was used, with emission detected between 600 nm and 680 nm.

Images were processed as maximum intensity projections of acquired Z-stacks using Zen Blue 2012 software. RACK1A-NUDT7 (Olejnik et al., 2011) and RACK1A-GPA1 (Cheng et al., 2015) pairs were used as positive and negative controls, respectively. The relative abundance of interaction was calculated as a ratio of the mean fluorescence intensity of YFP relative to mRFP. The experiment was performed twice, and at least one region of interest from 30 imaged cells per replicate was considered for calculations. The threshold ratio, accounting for background YFP fluorescence, was calculated from the analysis of negative control samples. Positive control was used to measure the ratio of known interactions, and the calculated average value was set to 1 for normalization. Statistical significance between studied interactions and negative control was evaluated using a one-way ANOVA test in GraphPad Prism 8.3.0.

### Bioinformatic predictions of PPIs

To predict possible direct PPI interface sites of RACK1A with FSD1, a bioinformatic approach utilizing AlphaFold 2 implementation based on ColabFold (Jumper et al., 2021; Mirdita et al., 2022), together with AlphaFold-Multimer (Evans et al., 2022) was performed. To run the predictions, https://neurosnap.ai/ (Neurosnap Inc. - Computational Biology Platform for Research. Wilmington, USA) interface was used according to the user’s manual. After modelling, 5 structures with highest predicted Local Distance Difference Test (pLDDT) and the highest combination of the predicted template modelling (pTM) and interface pTM (ipTM) scores (combined score = 0.8 ipTM + 0.2 pTM) (Homma et al., 2024) were extracted and uploaded to PDBsum (Laskowski, 2001) with the top one being used to create a scheme of possible amino acid residues in the PPI interface. SWISS-MODEL Workspace / GMQE (Waterhouse et al., 2018) Structure Assessment tool was used to generate models from obtained structures.

### Co-immunoprecipitation of RACK1A-GFP interacting proteins

The 14-day-old seedlings (2 g in fresh weight) of RACK1A-GFP line were treated with liquid ½ MS media with or without 150 mM NaCl for 1h and ground in liquid nitrogen. The protein extraction, co-immunoprecipitation and “on beads” trypsin digestion were performed as described in Hunter et al. (2019) and the peptides were cleaned on C18 cartridges (Bond Elut C18, 100mg, 1ml, Agilent Technologies). The LC-ESI-MS/MS analyses were performed on a nanoflow HPLC system (Easy-nLC1200, Thermo Scientific) coupled to the Q Exactive HF mass spectrometer (Thermo Scientific, Bremen, Germany). The peptide separation and data acquisition were performed as described in Hunter et al. (2019), while using Thermo Xcalibur 4.1 software (Thermo Fisher Scientific) for the data acquisition.

Raw data files were searched for protein identification using Proteome Discoverer 3.2 software (Thermo Scientific) connected to an in-house server running the Mascot 3.0.0 software (Matrix Science). Database search was performed against SwissProt database (version 2024_03) with taxonomy filter *Arabidopsis thaliana*. The experiment was performed in 4 biological replicates. Proteins identified by one peptide and proteins detected only in one biological replicate were excluded. Proteins interacting with YFP (Hunter et al., 2019) were considered unspecific and were removed.

### Whole Mount Immunolocalization

Four-day-old seedlings of WT and the transgenic RACK1A-GFP line were used for immunolocalization studies targeting TSN1/2 and RACK1A-GFP in root whole mounts following the protocol described by Šamajová et al. (2014). Prior to fixation, seedlings were incubated in liquid ½ MS medium or ½ MS medium supplemented with 100 mM NaCl for 30 min. Samples were immunolabeled overnight at 4°C with primary antibodies: anti-TSN1/2 rabbit polyclonal (diluted 1:75) and anti-GFP mouse monoclonal (diluted 1:100) in 3% (w/v) BSA in PBS. Subsequently, AlexaFluor 647 goat anti-rabbit and AlexaFluor 488 goat anti-mouse secondary antibodies (Invitrogen, USA) were applied at a 1:500 dilution in 2.5% (w/v) BSA in PBS for 2h at 37°C. Samples were observed by Zeiss LSM710 microscope (Carl Zeiss, Jena, Germany) equipped with Plan-Apochromat 40×/1.4 Oil DIC M27 objective. Fluorophore excitation was performed using lasers with lines at 488 and 631 nm. Image processing and analysis were carried out using Zeiss ZEN software (Black and Blue versions, Carl Zeiss, Germany), Photoshop 6.0/CS, and Microsoft PowerPoint. Quantitative colocalization analysis between RACKA1-GFP and TSN1/2 was performed on selected ROIs from single-plane confocal sections using the colocalization tool in Zeiss ZEN 2014 software (Black version).

## Results

### Microscopic analysis of tissue-specific and developmental RACK1A-GFP localization in Arabidopsis roots

Prior to examination of RACK1A-FSD1 interaction, we conducted a detailed tissue-specific and developmental localization of RACK1A-GFP in roots of *rack1a-1* mutant complemented with *proRACK1A::RACK1A:GFP* construct, to gain insights into possible synergies in functions of both proteins. Fluorescence signal of cellular RACK1A-GFP presence was observed in all plant organs of 5-day-old seedlings, while the strongest signal was observed in organs and tissues with high meristematic activity such as root tip, root primordia, and developing true leaves (Supplemental Figure 1).

In the root tip, RACK1A-GFP exhibited high abundance in meristematic zone (Supplemental Figure 2A). However, in cells of quiescent center, stem cell niche, columella and lateral root cap, the fluorescence intensity was substantially lower (Supplemental Figure 2A). On the subcellular level, in both in the root meristematic (Supplemental Figure 2B) and elongation (Supplemental Figure 2C, D) zones, a high abundance of RACK1A-GFP was observed in the cytosol, while in the nucleus, the fluorescence intensity was lower. ACLSM detection revealed that RACK1A may be involved in the lateral root initiation (Supplemental Figure 2E-H). Within the pericycle, RACK1A-GFP signal appeared substantially increased in lateral root founder cells, even before the first anticlinal cell division (Supplemental Figure 2E). Subsequently, high abundance of RACK1A-GFP was retained in newly formed cells after the first anticlinal cell division (Supplemental Figure 2F), which was also apparent during all subsequent phases of lateral root primordia development (Supplemental Figure 2G-H, Supplemental Video 1) as also indicated by pseudo-color-coding visualization (Supplemental Video 2).

In the root elongation zone, the abundance of RACK1A-GFP was significantly higher in trichoblasts compared to atrichoblasts (Supplemental Figure 2D), suggesting an involvement of RACK1A in root hair formation. During root hair growth, RACK1A-GFP was accumulated in the cortical cytoplasm of the bulging site at the basal end of trichoblasts (Figure 1A), in root hair bulges (Figure 1B) and actively growing root hairs (Figure 1C). At the subcellular level, RACK1A-GFP localization was abundant in nuclei and at the growing root hair tip (Figure 1C, D). Note that a similar pattern in growing root hairs was observed also for FSD1-GFP (Dvořák et al., 2021). Dynamic mode of developmental RACK1A-GFP localization during root hair formation was revealed by live-cell LSFM root imaging (Figure 1E, F; Supplemental Figure 3; Supplemental Video 3). Involvement of RACK1A in different stages of root hair development was documented by a tip-focused localization of RACK1A-GFP in bulges, emerging, and actively-growing root hairs. Importantly, it disappeared from the tip in fully-grown root hairs with terminated tip growth (Figure 1E; Supplemental videos 3, 4; Supplemental Figure 3A-F). Subcellular pattern of RACK1A-GFP fluorescence intensity distribution was confirmed by a pseudo-color-coding visualization (Figure 1F; Supplemental videos 5, 6; Supplemental Figure 3G-L) and by a kymograph-based semi-quantitative evaluation of growing root hairs (Figure 1G-J), evidencing a tip-focused RACK1A-GFP accumulation.

**Figure 1.**
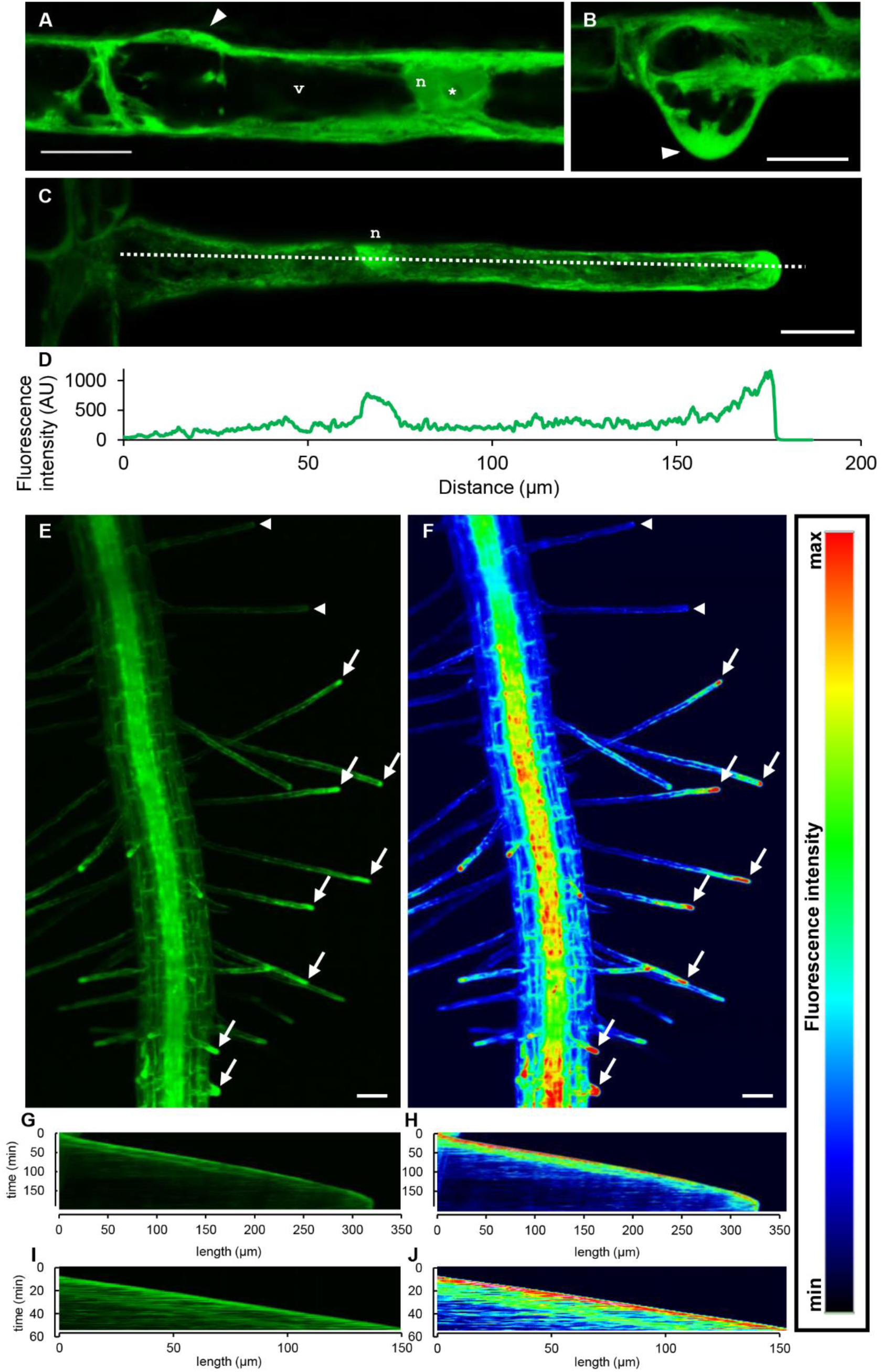
Microscopic observation of RACK1A-GFP subcellular localization during root hair development and tip growth. **(A-D)** RACK1A-GFP localization in a trichoblast at early (A) and late (B) bulging stage. Fluorescent RACK1A-GFP signal was located in a cytoplasm, particularly at the forming root hair tip (arrowheads in A, B) and in nucleus (n). Unlike that, nucleoli (*) and vacuoles (v) were devoid of RACK1A-GFP. In growing root hairs, RACK1A-GFP was located in cytoplasm, nucleus (n) and growing tip (C), as evidenced by a profile-based measurement of the fluorescence intensity distribution (D) along the line shown in (C). **(E, F)** *In vivo* dynamic localization of RACK1A-GFP in growing primary root visualized using light-sheet fluorescence microscopy in a root hair-formation zone. RACK1A-GFP (E) and pseudocolor (F) fluorescence intensity distribution at different stages of root hair development showed a tip-focused localization of RACK1A-GFP in bulges, emerging, and actively growing root hairs (arrows), while it disappeared from the tip in fully-grown root hairs with terminated tip growth (arrowheads). **(G-J)** Quantitative representation of root hair tip growth rate in kymographs of selected root hair until the termination of its tip growth (G, H), and depicted phase of its fast growth (I, J). Tip-focused fluorescence signal is presented as either RACK1A-GFP (G, I) or pseudocolor (H, J) fluorescence intensity distribution profile. Kymographs present measurements in 187 min and 350 µm (G, H), and 60 min and 150 µm (I, J) intervals. Heat map shows RACK1A-GFP fluorescence intensity in pseudocolors with the lowest fluorescence intensity corresponding to the dark blue and the highest fluorescence intensity corresponding to the red color. Scale bar = 20 µm (A-C) and 50 µm (E, F).

### RACK1A interacts with FSD1, predominantly in the cytosol

Published high-throughput yeast two-hybrid screening identified FSD1 as a potential interactor of RACK1A (Guo et al., 2019). To discover more about the possible RACK1A-FSD1s interactions regarding the PPI interface, a bioinformatic approaches based on AlphaFold2 and AlphaFold-Multimer were employed.

As shown in Figure 2A, B, the first of five possible predicted complexes with the best combined pTM and ipTM scores was extracted and modelled with SWISS-MODEL Workspace / GMQE Structure Assessment tool. The pLDDT score of analyzed complex was 88.83 and represents very high accuracy of each predicted residue’s spatial orientation and position by AlphaFold2. pTM score corresponding to overall topological accuracy was 0.64. The highest obtained ipTM representing the degree of reliability of the predicted interaction interfaces was 0.2. Combined pTM and ipTM scores, representing the overall PPI prediction accuracy, were 0.29.

**Figure 2.**
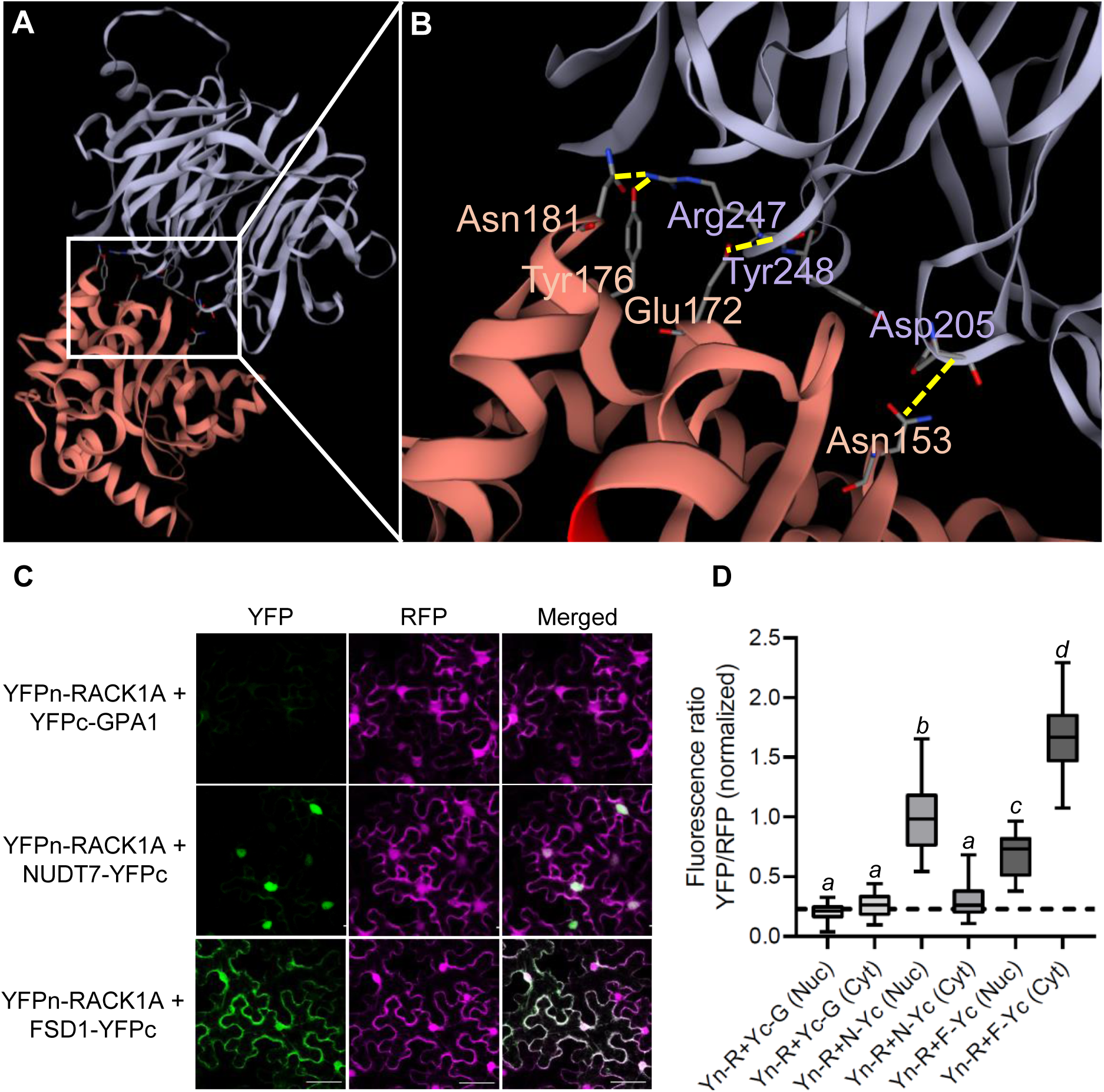
Functional validation of FSD1-RACK1A interaction. **(A, B)** Bioinformatic prediction of direct protein-protein interaction (PPI) interface between RACK1A and FSD1. (A) Model of RACK1A-FSD1 interaction. White frames indicate the predicted PPI interfaces. (B) Details of predicted PPI interfaces. Amino acid residues that are predicted to be involved in hydrogen bonds are interconnected by yellow dashed line. https://neurosnap.ai/ interface utilizing AlphaFold 2 implementation based on ColabFold, and AlphaFold-Multimer were used to run the PPI interface predictions. SWISS-MODEL Workspace / GMQE Structure Assessment tool was used to generate models from obtained structures. **(C, D)** Protein-protein interaction (PPI) study of RACK1A with FSD1 by ratiometric bimolecular fluorescence complementation assay (rBiFC). (C) Representative images of rBiFC assay taken by confocal laser scanning microscopy (CLSM). YFP signal represents the PPI, RFP signal functions as a control of transformation and reference for calculations of relative PPI strength. Scale bar = 20 µm. (D) Relative quantification of PPI strength measured as a ratio of YFP to RFP relative fluorescence intensity in nuclei (Nuc) or cytosol (Cyt). Ratio values were normalized to the mean ratio measured in nuclei of positive control transformed with vectors carrying *YFPn-RACK1A+NUDT7-YFPc* (Yn-R+N-Yc) genes. The dashed line represents the mean ratio measured in negative controls transformed with vectors carrying YFPn-RACK1A + YFPc-GPA1 (Yn-R+Yc-G) and functions as a threshold of positive interaction. Vector carrying *YFPn-RACK1A+FSD1-YFPc* (Yn-R+F-Yc) genes was used to study the interaction of RACK1A with FSD1. Italic letters indicate a statistically significant difference at a p < 0.05 as determined by one-way ANOVA with post-hoc Tukey HSD test.

To uncover the possible inter-molecular interactions of RACK1A with FSD1, obtained AlphaFold2 data were analyzed using PDBsum. In RACK1A-FSD1 complex, 12 residues of RACK1A covering 587 Å^2^ interface area and 9 residues of FSD1 covering 686 Å^2^ interface area were predicted to be directly involved in the interaction (Figure 2A, B). Of these residues, Asp205, Arg247 and Tyr248 in RACK1A and Asn153, Glu172, Tyr176 and Asn181 in FSD1 were predicted to form 5 hydrogen bonds in the interface (Figure 2B). Of note, these FSD1 residues are situated in proximity to Asp169 and His173, the Fe^3+^-binding residues essential for FSD1 activity (Pilon et al., 2011). Moreover, up to 155 non-bonded contacts were predicted between all interface residues. Notably, all mentioned residues in RACK1A, together with Glu172 and Asn153 in FSD1, were also predicted in the second-ranked model (data not shown).

Since both proteins participate in plant salt stress responses, we analyzed the salt-induced interactome of RACK1A-GFP in a stably transformed Arabidopsis line using GFP trap technology coupled with MS/MS (Supplemental Tables 3 and 4). This analysis revealed that a substantial portion of the RACK1A-GFP interactome was salt stress-dependent, as evidenced by the higher number of protein interaction clusters identified by STRING compared to the interactome of the mock treated control plants (Supplemental Figures 4 and 5). The salt-induced RACK1A interactome was enriched with proteins involved in ribosome biogenesis, Golgi vesicle transport, metabolism and gene expression (Supplemental Figure 6). Gene ontology annotations uniquely induced by salt stress included Golgi vesicle transport, amino acid metabolism, and cell tip growth (Supplemental Tables 5 and 6). Notably, the interactome also included proteins associated with oxidative stress responses, such as FSD1, FSD2, FSD3, MANGANESE SOD 1 (MSD1), ASCORBATE PEROXIDASE 3, secretory peroxidases, and peroxiredoxins (Supplemental Table 4). These findings further validate the interaction between RACK1A and FSD1.

Our next objective was to validate the interaction between RACK1A and FSD1 and to determine their subcellular localization using a ratiometric BiFC assay, allowing semi-quantitative analysis of PPIs *in vivo* (Figure 2C, D). The presence of RFP fluorescence confirmed the successful transient expression of all expression vectors containing the genes of interest in the leaf epidermal cells of *N. benthamiana* (Figure 2C). The lack of YFP fluorescence in samples expressing the negative control (RACK1A-GPA1) and the presence of YFP fluorescence in samples with the positive control (RACK1A-NUDT7) validated the functionality of the method. The interaction between RACK1A and FSD1 was predominantly localized in the cytosol (Figure 2C), with the ratio of YFP to RFP fluorescence intensity being higher than that observed in the positive control measurements (Figure 2D). Notably, the abundance of nuclear YFP signal was significantly lower compared to the signal measured in the positive control, indicating that the RACK1A-FSD1 interaction primarily occurs in the cytosol and is considerably less abundant in the nucleus.

### SOD activities and abundances in rack1a mutants and fsd1-1 rack1a-1 double mutant

To study genetic interaction between RACK1A and FSD1, a double mutant lacking both *FSD1* and *RACK1A* was generated through conventional crossing, and the absence of these proteins was confirmed via immunoblotting (Supplemental Figure 7A, B). Functional genetic experiments also included single *fsd1-1* and *rack1a-1* mutants, as well as newly developed CRISPR/Cas9 knock-out mutant *rack1a-5*, all lacking corresponding proteins (Supplemental Figure 7A, B).

Abundances and activities of SOD isozymes were evaluated in plants grown under limited Cu^2+^ availability (0.05 µM) (Figure 3) causing increased *FSD1* expression, while minimizing the impact of *CSD1*, which is weakly expressed (Melicher et al., 2022). As expected, the in-gel SOD activity assay revealed that FSD1 exerted the highest activity among the SOD isozymes in WT (Figure 3A, B). On the other hand, the activities of SOD isozymes in both *rack1a* mutants were significantly higher compared to WT (Figure 3A, B). The activity of MSD1 was 1.7 and 1.6-times higher in *rack1a-1* and *rack1a-5* mutants, respectively. Moreover, the activities of FSD1 and both CSDs were 1.55 and 2-times higher, respectively, in *rack1a-1* mutant compared to WT. Comparably, in *rack1a-5* mutant, FSD1 activity was 1.54-times higher, while CSDs activity was increased up to 2.18-times. These data suggest the regulation of SOD isozymes activities by RACK1A.

**Figure 3.**
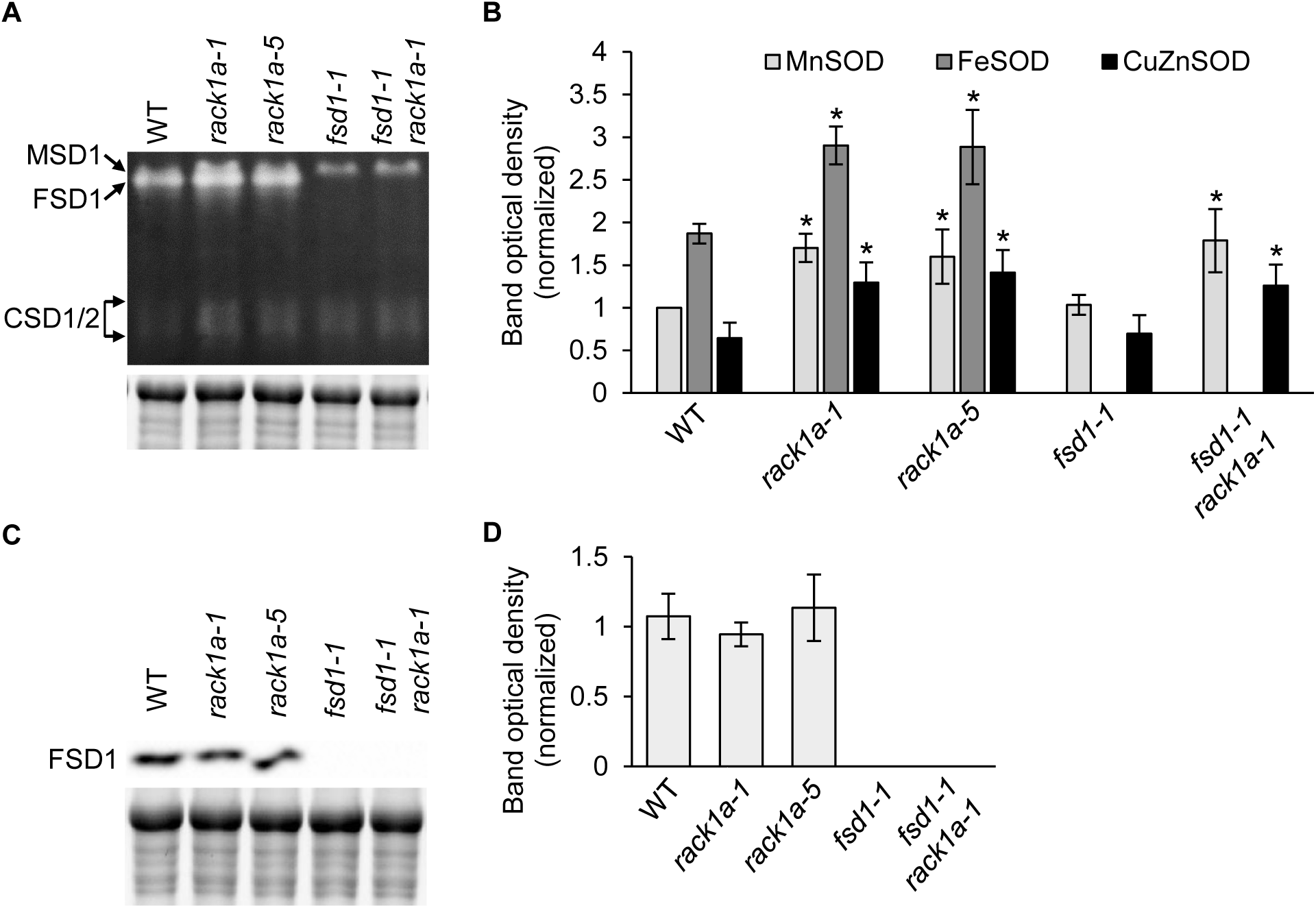
In-gel superoxide dismutase (SOD) isozyme activity measurement and immunoblotting of FSD1 in 14-day-old wild type (WT), *rack1a-1*, *rack1a-5*, *fsd1-1* mutant lines and *fsd1-1 rack1a-1* double mutant grown on ½ MS media. **(A)** In-gel SOD activity staining supplemented with respective controls of protein loading using Stain-free gel. **(B)** Quantification of band optical density in A. Values in B are expressed as relative to the activity of MSD1 (mean ± SD, N = 4). **(C)** Immunoblots of Fe-superoxide dismutase (FSD1), using anti-FSD1 antibody, supplemented with respective controls of protein loading using Stain-free gels. **(D)** Quantification of band optical density in C. Values in D are expressed as relative to the abundance of the protein in WT (mean ± SD, N = 4). Asterisks indicate a statistically significant difference between WT and respective line as revealed by one-way ANOVA with post-hoc Tukey HSD test (* indicates statistical significance at p < 0.05).

Immunoblotting showed that in both *rack1a* mutants, the abundances of FSD1 were similar to WT (Figure 3C, D). Thus, no positive correlation between FSD1 abundance and activity was observed, suggesting the regulation of FSD1 activity by RACK1A, possibly by the PPI.

### Both single rack1a mutants and fsd1-1 rack1a-1 double mutant have altered phenotypes

Phenotypic analysis of *rack1a-1* lines complemented by RACK1A-GFP*, rack1a-1, rack1a-5, fsd1-1* single and *fsd1-1 rack1a-1* double mutants showed genetic interaction between FSD1 and RACK1A affecting plant development (Figure 4A-C). Measurement of fresh weight of 10-day-old seedlings revealed a significant decrease in both *rack1a* mutants as well as in *fsd1-1 rack1a-1* double mutant (Figure 4B). While WT, RACK1A-GFP and *fsd1-1* mutants exhibited the highest fresh weights 10 days after germination, the fresh weights of both *rack1a* mutants were reduced. The fresh weight of *fsd1-1 rack1a-1* mutant was the lowest among the tested lines (Figure 4B).

**Figure 4.**
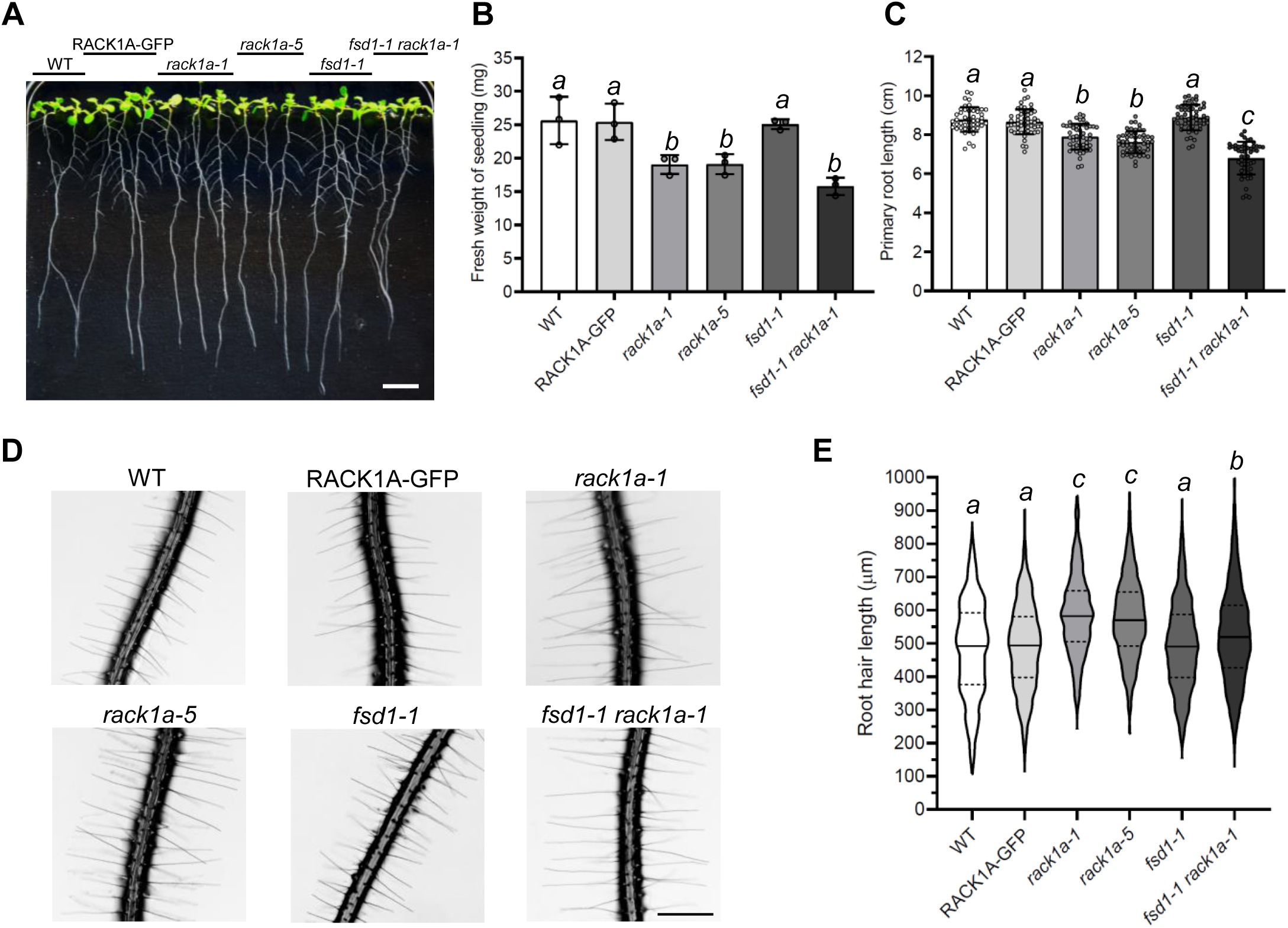
Phenotypic analysis of Col-0 (wild type; WT), RACK1A-GFP-complemented *rack1a-1* mutant, *rack1a-1*, *rack1a-5, fsd1-1* mutants, and *fsd1-1 rack1a-1* double mutant. **(A)** Representative picture of 10-day-old seedlings. **(B, C)** Quantification of fresh weight (B) and primary root length (C) of 10-day-old seedlings. Phenotypic analyses were performed in three repetitions (N = 90 for fresh weight, N = 50 for primary root length). Circle symbols represent average value from 30 seedlings in B or individual values in C. Error bars represent standard deviation. Italic letters indicate a statistically significant difference at a p < 0.05 as determined by ordinary one-way ANOVA test. Scale bar = 1 cm. **(D, E)** Phenotypic analysis of root hair length in Col-0 (wild type; WT), RACK1A-GFP-complemented *rack1a-1* mutant, *rack1a-1*, *rack1a-5, fsd1-1* mutants, and *fsd1-1 rack1a-1* double mutant. (D) Representative pictures of 7-day-old seedlings mature root hairs measured at a distance of more than 5 mm from the root tip. (E) Quantification of root hair length. Phenotypic analysis was performed in three repetitions (N = 1200). Violin plots represent the distribution of individual root hair length values. Lines represent the median value, and dashed lines represent quartile values. Different italic letters above columns indicate a statistically significant difference at a p < 0.05 as determined by Kruskal-Wallis One Way Analysis of Variance on Ranks. Scale bar = 500 µm.

Similarly to fresh weight, the primary root length of 10-day-old WT, RACK1A-GFP and *fsd1-1* plants exhibited similar values. Conversely, *rack1a-1* and *rack1a-5* mutants were shorter in root length, and *fsd1-1 rack1a-1* double mutants exhibited the shortest primary roots (Figure 4C). Therefore, *FSD1* and *RACK1A* show additive genetic influence in regulation of fresh weight and primary root length.

Both proteins exhibit accumulation in tips of growing root hairs, indicating possible genetic interaction in root hair growth regulation. To investigate this, we analyzed root hair length in all studied lines (Figure 4D, E). In WT, the root hair length median value was 492 µm, with interquartile ranging from 376 µm to 591 µm. Comparably, the median values of root hair length in complemented line and *fsd1-1* mutant were 493 µm and 490 µm, respectively. Quartile values ranged from 398 µm to 580 µm in complemented line and from 397 µm to 586 µm in *fsd1-1* mutant, showing high similarity when compared to WT. On the contrary, root hair length in both *rack1a* mutants, as well as in *fsd1-1 rack1a-1* double mutant was significantly higher. While in *rack1a-1* mutant and *rack1a-5* mutant, the median values reached 582 µm (with quartiles ranging from 505 µm to 658 µm) and 570 µm (with quartiles ranging from 492 µm to 654 µm), respectively, in *fsd1-1 rack1a-1* double mutant, the median value was substantially lower with 519 µm. Additionally, the root hair length of *fsd1-1 rack1a-1* double mutant was significantly higher than the length measured in WT, complemented line and *fsd1-1* line, thus displaying an intermediate phenotype between *fsd1-1* and *rack1a-1* mutant (Figure 4D, E). The results suggest a genetic interaction between FSD1 and RACK1A, involved in regulation of root hair growth and primary root development.

### Subcellular relocation of RACK1A in response to salt stress

Despite the roles of RACK1 in plant stress responses are intensively studied, limited attention is devoted to the conditional dynamic relocation of this protein. Therefore, we aimed to determine the changes in localization and intensity of the RACK1A-GFP signal in response to salt stress (Figure 5). The meristematic cells of the root tip were imaged immediately after perfusion of the liquid media (t = 0 min) and again after 30 minutes. In control conditions, no change in the intensity or localization of the RACK1A-GFP signal was observed during the observation (Figure 5A). Likewise, no changes were observed immediately after the addition of 100 mM NaCl (t = 0 min; Figure 5A). On the other hand, incubation of seedlings with 100 mM NaCl for 30 min caused the formation of small, strongly fluorescent dynamic condensates, resembling stress granules (SG). These granular structures varied in size and were localized in the cytosol but not in the nucleus (Figure 5A). Recovery from the salt stress by exchanging the NaCl-containing medium with control ½ MS medium resulted in the immediate disappearance of these SG-like structures (Figure 5B). We also tested their formation under simultaneous effect of NaCl and SG inhibitor cycloheximide. Importantly, the presence of cycloheximide in the NaCl-containing medium prevented the formation of these structures, supporting the hypothesis about their identity (Figure 5C).

**Figure 5.**
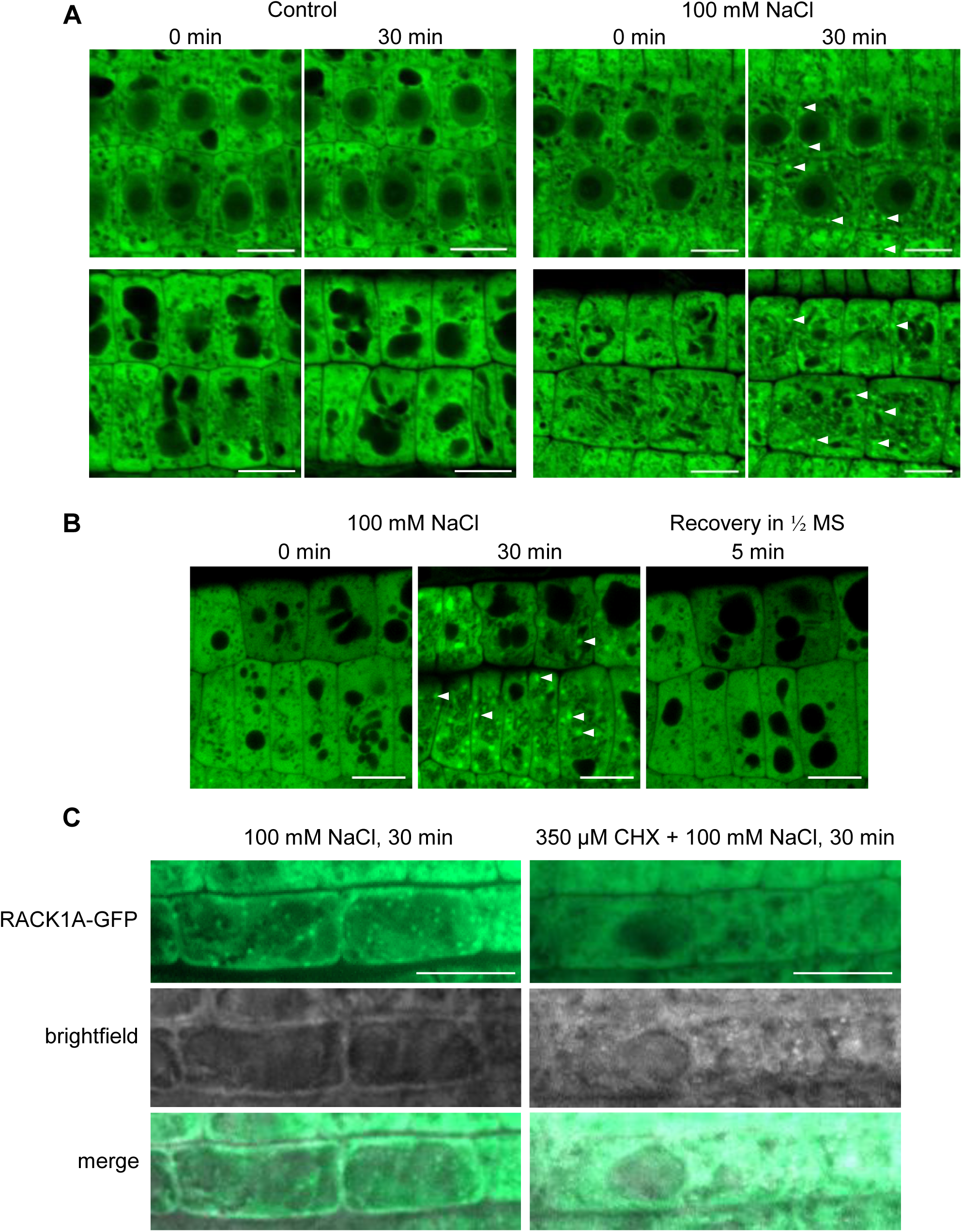
RACK1A-GFP localization in root meristematic zone cells upon 100 mM salt stress. **(A)** Root meristem cells before and 30 min after the treatment with ½ MS (Control) and 100 mM NaCl. Images were taken from middle focal plane containing nucleus and cytosol (upper part of the images) or upper focal plane containing cytosol near plasma membrane (bottom of the pictures). White arrowheads indicate a condensation of the fluorescence signal. Scale bar = 10 µm. **(B)** RACK1A-GFP localization in root meristematic zone cells after replacement of NaCl with liquid ½ MS medium. Images were taken from cortical focal plane. White arrowheads indicate a condensation of the fluorescence signal. Scale bar = 10 µm. **(C)** RACK1A-GFP localization in root epidermal cells after the treatment with 100 mM NaCl and 100 mM NaCl with 350 µM cycloheximide (CHX). Scale bar = 20 µm.

We examined the subcellular distribution of RACK1A-GFP and TSN proteins (TSN1 and TSN2) in whole-mount root samples of *rack1a-1* mutants complemented with RACK1A-GFP by immunolocalization. Under control conditions, RACK1A and TSN proteins were predominantly localized in the cytoplasm, with partial colocalization and association (Supplemental Figure 8A). However, upon exposure to salt stress, both proteins partially relocalized to SG (Supplemental Figure 8B). To quantify the extent of colocalization between RACK1A and TSN proteins, we performed a quantitative colocalization analysis, which revealed a higher degree of colocalization following salt stress (Supplemental Figure 8C-E).

### Colocalization of RACK1A with FSD1

Next, we examined the colocalization of FSD1-mRFP and RACK1A-GFP fusion proteins in a rescued *rack1a* mutant line. In root epidermal cells, both proteins colocalized in cortical cytoplasm and, upon 30 minutes of salt treatment, showed visual colocalization in structural condensates resembling stress granules (Figure 6A, B). Notably, cycloheximide treatment prevented this colocalization, as well as the formation of structural condensates (Figure 6C, D). Colocalization of RACK1A-GFP with FSD1-mRFP in structural condensates was confirmed by qualitative (Figure 6E-I) and semi-qualitative (Figure 6J-M) colocalization analyses.

**Figure 6.**
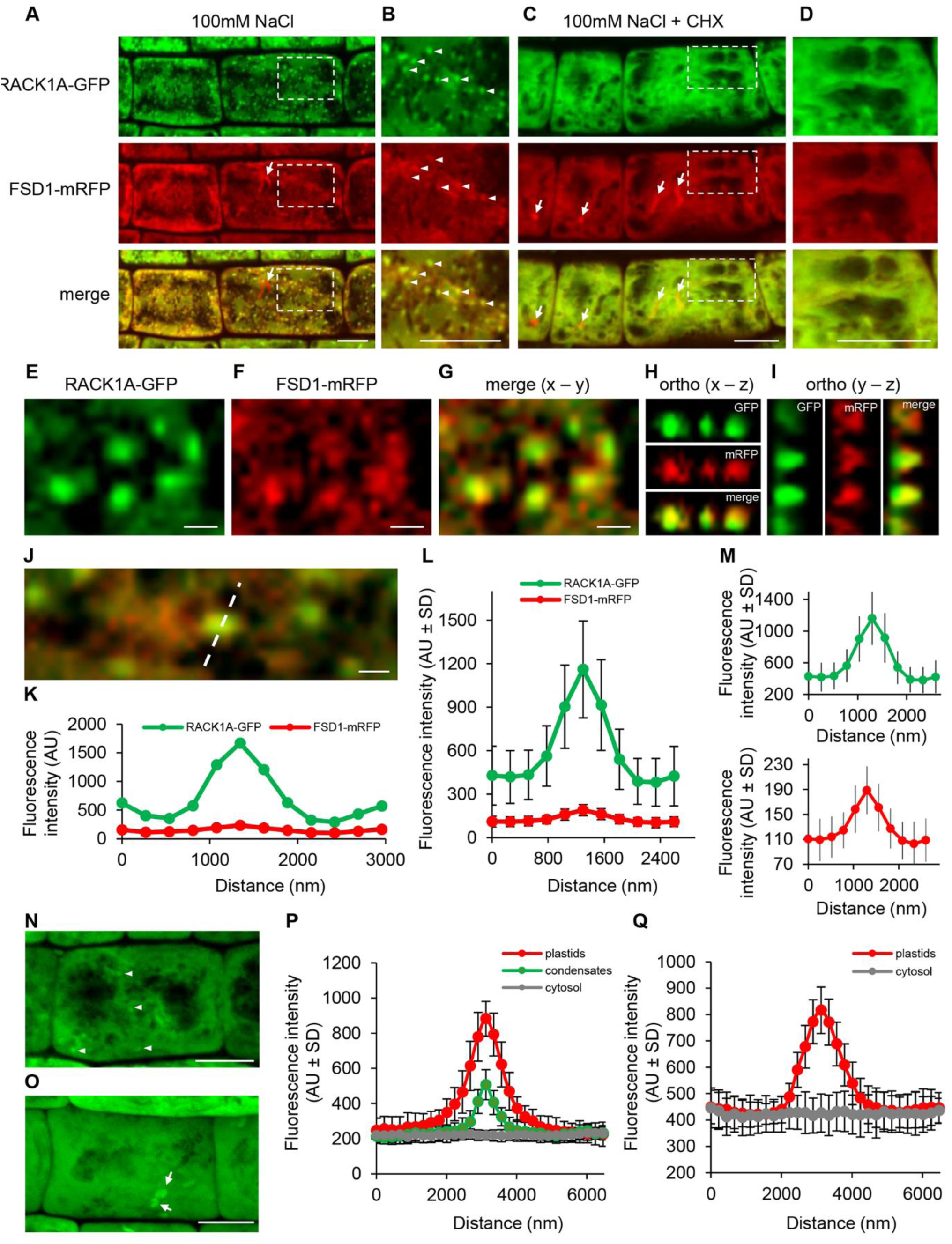
Subcellular localization analysis of RACK1A-GFP and FSD1-mRFP in root epidermal cells of the *rack1a-1* mutant after NaCl and cycloheximide treatments. **(A-D)** Overview (A, C) and detailed view (B, D) of cortical cytoplasm in epidermal cells of the root transition zone of the genetically-rescued *rack1a-1* mutant showing distribution of RACK1A-GFP (green), FSD1-mRFP (red) and their overlay (merge) after treatment with ½ MS containing 100 mM NaCl (A, B) or 100 mM NaCl + 350 µM cycloheximide (CHX; C, D) for 30 min. The detailed views (B, D) show close-ups marked by the dashed rectangle insets in (A, C). White arrowheads indicate structural condensates accumulating both GFP and mRFP fluorescence signals. **(E-I)** Qualitative colocalization analysis after treatment with ½ MS containing 100 mM NaCl for 30 min between RACK1A-GFP (E) and FSD1-mRFP (F) in an overlay image in a x – y view (G) and an orthogonal x – z (H) and y – z (I) spatial views in structural condensates from marked cell in (A). **(J-M)** Semi-quantitative colocalization analysis between RACK1A-GFP and FSD1-mRFP in a selected structural condensate (J) after treatment with ½ MS containing 100 mM NaCl for 30 min by analysis of fluorescence intensity (K) along the profile marked by a white line in (J). Averaged fluorescence intensity distribution from a profile measurement of structural condensates (n = 49) showed colocalization of GFP and mRFP fluorescence peaks (L), substantiated after their separate display with relevant range of fluorescence intensity distribution values (M). Arrows in (A, C) point FSD1-mRFP localized in plastids. (N-Q) Semi-quantitative analysis of FSD1-GFP fluorescence signal distribution after 100 mM NaCl treatment (n = 20) in plastids, condensates, and cytosol of *fsd1* mutant (N, P) and in plastids and cytosol of *rack1-1* mutant (O, Q). White arrowheads indicate structural condensates and arrows indicate plastids. Scale bar = 20 µm (A-D, N, O), 1 µm (E-G, J).

### FSD1-GFP localization in rack1a mutant

To reveal putative RACK1A-mediated or RACK1A-controlled alterations in FSD1 subcellular localization after salt stress, we generated stably transformed *rack1a* lines expressing FSD1-GFP under its native promoter. The localization pattern of FSD1-GFP in *rack1a-1* mutant was compared with the complemented *fsd1-1* mutant (Dvořák et al., 2021). The observation of FSD1-GFP in *fsd1* and *rack1-1* mutants showed that while the signal of FSD1-GFP in *fsd1-1* mutant, in addition to its accumulation in plastids, was homogenously distributed in cortical cytoplasm in control conditions and FSD1-GFP accumulated in structural condensates upon 30 minutes salt treatment (Figure 6N, P). Such relocation and accumulation of FSD1-GFP in structural condensates did not occur in *rack1a* mutant (Figure 6O, Q). Therefore, this genetic evidence suggests that RACK1A is indispensable for the condensate-specific FSD1 localization upon salt stress.

### ROS levels are differently modulated in mutant and transgenic lines upon salt stress response

Previously, we determined the salt stress susceptibility of *fsd1* mutants, showing less viable plants after salt treatment compared to WT (Dvořák et al., 2021). Here, results indicate an increased salt stress resistance of the single *rack1a-1* and *rack1a-5* mutants. The simultaneous absence of FSD1 and RACK1A leads to a salt stress response intermediate between WT and the single mutants (Figure 7A, B). These data indicate that *FSD1* is epistatic to *RACK1A* in salt stress response, and both genes jointly modulate the salt stress responses in Arabidopsis. Finally, we also examined the intracellular ROS levels in root cells with the vital fluorescent dye CellRox Deep Red, which is labelling superoxide and hydroxyl radicals. As shown previously, the hypersensitive *fsd1* single mutants showed higher fluorescence compared to WT in response to salt stress, indicating more intensive ROS accumulation in root cells (Dvořák et al., 2021). This vital histochemical staining showed significantly weaker fluorescence in roots of *rack1a* mutants as compared to WT (Figure 7C, D). The *fsd1 rack1a1* double mutant showed intermediate ROS fluorescence in response to salt (Figure 7C, D), while statistically significant difference was accounted between the single and double mutants. Collectively, the presented data support the regulatory role of RACK1-FSD module in Arabidopsis salt stress responses.

**Figure 7.**
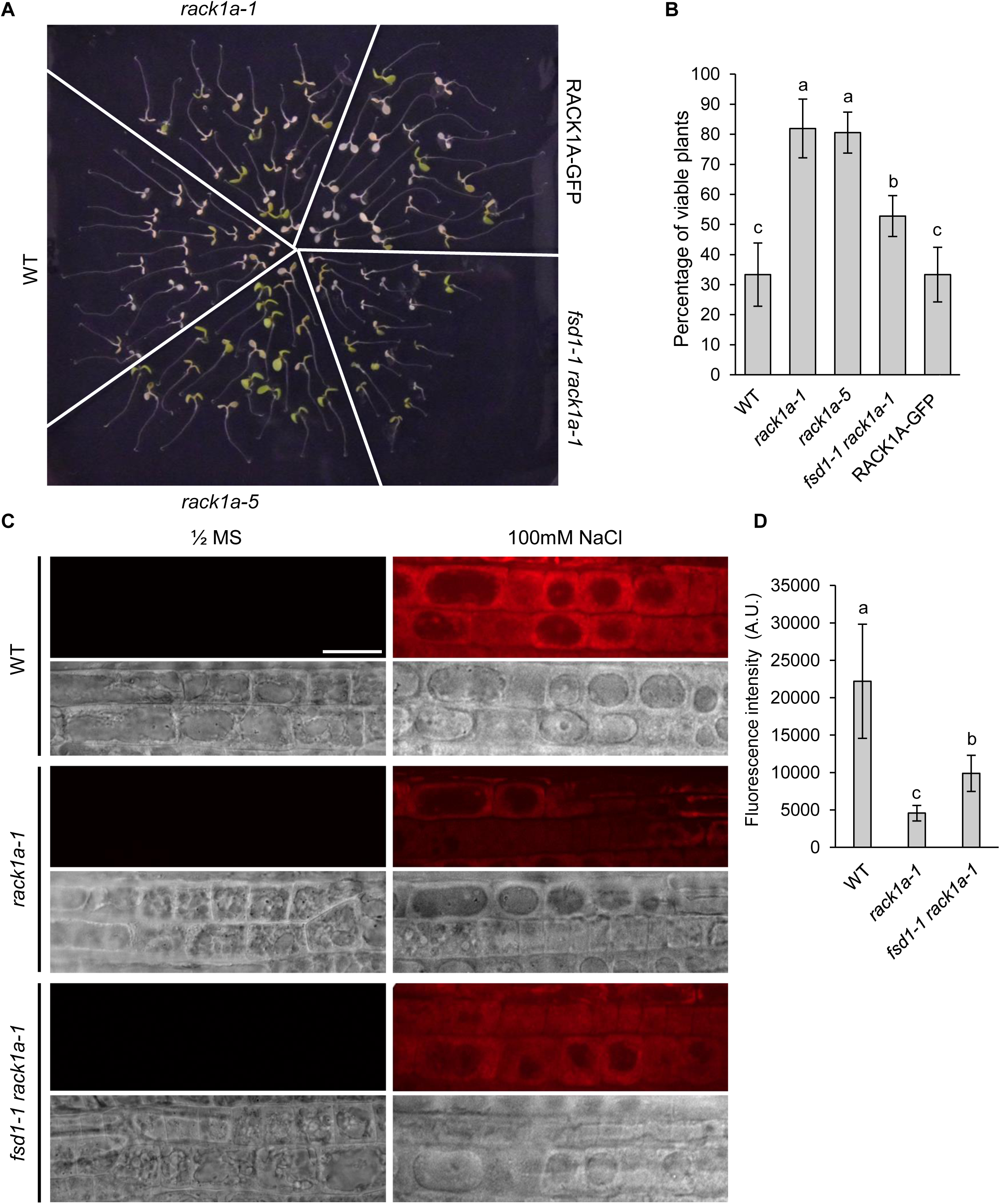
Analysis of NaCl-induced oxidative stress response in Col-0 (wild type; WT), *rack1a-1*, *rack1a-5* and *fsd1-1 rack1a-1* mutants, and RACK1A-GFP line. Five-day-old seedlings were transferred to a NaCl-containing medium. **(A)** Representative images of seedlings on the fifth day after the transfer to 150 mM NaCl-containing medium. **(B)** Quantification of seedlings with affected viability from (A). Plants with fully bleached cotyledons were considered unviable. Analysis was performed in three repetitions (N = 70). Different italic letters above the columns indicate a statistically significant difference between the respective lines and treatments at a p < 0.05 as determined by ordinary one-way ANOVA test. **(C)** ROS distribution visualized by fluorescent tracker CellRox Deep Red reagent in plasmolyzed root epidermal cells of Col-0 (wild type; WT), *rack1a-1*, *fsd1-1 rack1a-1* lines. ROS accumulation in mock-treated (½ MS) and plasmolyzed root epidermal cells (½ MS with 100 mM NaCl) visualized by fluorescent tracker CellRox Deep Red reagent. The observed regions of root epidermal cells were visualized using transmitted light. Scale bar = 20µm **(D)** Quantification and statistical evaluation of CellRox Deep Red reagent fluorescence intensity (N = 15) as determined by one-way ANOVA with post-hoc Tukey HSD test.

## Discussion

Previous studies suggested the involvement of RACK1 in salt stress responses with distinct roles in different plant species (Guo et al., 2009; Li et al., 2017; Zhang et al., 2018; Denver and Ullah, 2019). However, the subcellular localization of RACK1A upon salt stress was not reported so far. In this study, we observed that RACK1A-GFP localizes to SGs upon salt stress. These are membrane-less cytoplasmic condensates of proteins, RNAs and metabolites that are formed after stress treatments (Maruri-López et al., 2021). They facilitate the plant stress responses by inhibiting translation, mediating mRNA turnover and protecting proteins from unfolding or degradation (Vanderweyde et al., 2013; Protter and Parker, 2016). Previous study has shown that SG marker protein TUDOR-SN PROTEIN 1 (TSN1) localizes to SGs upon salt stress, and *TSN1* RNAi lines exhibit higher salt stress sensitivity (Yan et al., 2014). SGs contain also other salt stress-related proteins such as OLIGOURIDILATE BINDING PROTEIN 1b (UBP1b) (McCue et al., 2012) and ANGUSTIFOLIA (Bhasin and Hülskamp, 2017). Mammalian RACK1 was identified as a new component of SGs, and its sequestration to these granules determines the cell fate upon diverse stress stimuli (Park et al., 2020; Masi et al., 2022). Nevertheless, the accumulation of plant RACK1A in SGs was not reported so far.

Our study uncovered an entirely new, SG-mediated mode of antioxidant enzyme regulation because it shows that FSD1, an important superoxide decomposing enzyme, relocalizes to SGs during salt stress. Notably, previous proteomic studies proposed the SG accumulation of FSD1 (Kosmacz et al., 2019). According to our study, RACK1A may recruit FSD1 to SGs, and its release from these condensates may facilitate salt stress tolerance. Further investigations will determine whether RACK1A may participate in the coordination of superoxide and H_2_O_2_ scavenging machinery in plants. The above-mentioned proteomic analysis also proposed a SG localization of H_2_O_2_ decomposing enzyme ASCORBATE PEROXIDASE 1 (APX1), which is a potential RACK1A interactor (Guo et al., 2019). The immunoblotting and in-gel SOD activity assays revealed that RACK1A negatively regulates FSD1 activity, suggesting RACK1A as a possible new FSD1 activity repressor. Previous studies reported that interactions of RACK1A with various proteins alter their subcellular localization and function (Wang et al., 2019; Li et al., 2023; Masood et al., 2023; Fu et al., 2024; Li et al., 2024). Analysis of PPI interface implicated in RACK1A-FSD1 interaction by AlphaFold-Multimer revealed that Tyr248, a RACK1A phosphorylation site crucial for PPIs and RACK1A homodimerization (Kundu et al., 2013, Sabila et al., 2016), may participate in the interaction. RACK1A homodimerization positively affects tolerance to UV-B induced oxidative stress in yeast (Sabila et al., 2016). According to the first and the second models predicted by Alphafold multimer, Tyr248, Arg247 and Asp205 in RACK1A may form hydrogen bonds or salt bridges with Glu172, Tyr176 and Asn153 in FSD1. All three mentioned FSD1 residues are situated in proximity to Asp169 and His173, the Fe^3+^-binding residues essential for FSD1 activity (Pilon et al., 2011). Thus, it is possible that via interaction, RACK1A may prevent FSD1 activation by masking the Fe^3+^ incorporation to the active site or affect the catalytic efficiency. Moreover, Glu172 and Tyr176 of Arabidopsis FSD1, proposed by our models as masked by RACK1A via interaction, are homologous to Glu198 and Tyr202 of *Vigna unguiculata* FeSOD, respectively (Supplemental Figure 9). Both amino acid residues were proposed to be important for the activity of the neighboring FeSOD monomer (Supplemental Figure 9; Muñoz et al., 2005). Disruption of this homodimer interaction interface results in up to 40% loss of activity as based on the studies of *E. coli* MnSOD (Edwards et al., 2001). Thus, RACK1A may suppress the activity of FSD1 by hindering its ability to dimerize. Taking together, apart from CPN20 (Kuo et al., 2013) and WRKY53 (Andrade Galan et al., 2024), also RACK1A may serve as a novel regulator of FSD1 activity.

We have found that RACK1A functions jointly with FSD1 to regulate specific developmental programs, such as root hair tip growth. Root hair development consists of a bulge formation and subsequent polarization of the cytoskeleton, membrane trafficking, and cell wall deposition at the tip to ensure polar expansion and root hair growth (Šamaj et al., 2006; Kuběnová et al., 2022). ROS are essential factors participating in root hair elongation (Schoenaers et al., 2017; Kuběnová et al., 2023). They are generated by RBOH family of enzymes (Torres et al., 2002; Sagi, 2006; Kuběnová et al., 2022) at the plasma membrane and released to apoplast. Root hairs, however, shows also prominent cytoplasmic and mitochondrial accumulation of ROS (Kuběnová et al., 2023). In this study, we elucidated an involvement of RACK1A in root hair growth and development, as *rack1a* mutants exhibited significantly longer root hairs when compared to WT. Moreover, RACK1A-GFP accumulated in the tip of growing root hairs, similar to FSD1-GFP, as reported previously (Dvořák et al. 2021). Interestingly, rice RACK1B interacts with N-terminus of RBOHD and overexpression of corresponding *OsRACK1B* gene results in H_2_O_2_ overaccumulation in pollen, which is negatively affecting elongation of pollen tubes (Rahman et al., 2022). Notably, knock-out double mutant in *RACK1A* and *FSD1* showed partial rescue of longer root hair phenotype typical for *rack1a* single mutants. Therefore, FSD1 might be an important factor modulating root hair elongation in the RACK1A absence. Since the RACK1A-FSD1 interaction occurs in the cytosol, and FSD1 does not localize to apoplast, RACK1A likely alters the FSD1 activity to modify cytosolic ROS during root hair elongation.

In summary, we identified RACK1A as a SG localized protein upon salt treatment. It sequesters FSD1 to SG during salt stress and alters its activity. The absence of RACK1A leads to FSD1 activation, prevention of its relocation to SGs and higher salt stress resistance of seedlings. The RACK1A-FSD1 module is also involved in developmental programs such as root hair elongation. These results reveal a novel mechanism of salt stress response in Arabidopsis.

## Acknowledgements

We thank Hemayet Ullah for providing the *rack1a-1* mutant and Michaela Tichá for help with the design of CRISPR/Cas9 mediated mutagenesis experiment. We also thank Katrin Janik for providing the entry vectors for rBiFC. Mass spectrometry analyses were performed at the Turku Proteomics Facility supported by Biocenter Finland.

## Author contributions

P.M., P.D., T.T., M.T., J.Ř., K.H., and O.Š. performed the experiments and analyses. T.T. and P.D. also assisted with data assessment. J.Š. provided the infrastructure. P.M., P.D., and T.T. drafted the manuscript, which was revised and edited by T.T., M.O. and J.Š. P.D. secured the funding. T.T. and P.D. conceived and supervised this study. All authors approved the final version of the manuscript.

